# Natural variations in reproductive timing of brood size and X-chromosome nondisjunction in *Caenorhabditis elegans*

**DOI:** 10.1101/2021.04.08.439028

**Authors:** Jiseon Lim, Jun Kim, Junho Lee

## Abstract

Reproductive timing underlies self-sperm production and X-chromosome nondisjunction rate in *Caenorhabditis elegans*. These traits can be measured as brood size and male-production rate, and brood size decreases and male-production rate increases as worms age. These phenotypic changes occur simultaneously; however, whether these changes share common underlying genetic architectures still remains unclear. To enhance our understanding of reproductive timing in *C. elegans*, we measured and compared these two reproductive timing-related traits of 96 wild strains during early, late and total reproductive periods. We found that the two traits exhibited natural phenotypic variation with few outliers and a similar reproductive timing pattern as previous reports. The brood size and the male-production rate were not genetically correlated, implying that they might have different genetic architectures and that meiosis quality can be maintained despite increased progeny-production in the late reproductive period caused by more self-sperm. We also identified loci and candidate genetic variants significantly associated with male-production rate in the late and total reproductive periods. Our results provide an insight into life history traits in wild *C. elegans* strains.

## Introduction

Reproductive timing is a temporarily regulated pattern of reproduction. In a selfing androdioecious nematode *Caenorhabditis elegans*, progeny-production rate of unmated hermaphrodites is the highest on the second day of adulthood and dramatically declines thereafter, whereas adult worms mostly die at day 15–20. Thus, these individuals produce approximately 90% of their progeny during the first 3 days of adulthood (Klass 1977; Huang et al. 2004; Zhang et al. 2021). This transient reproduction is a result of the sperm-limited fecundity of protandrous *C. elegans*. In *C. elegans*, spermatogenesis occurs during the last larval stage, and a shared pool of germ-cell precursors turns irreversibly into oogenesis after the completion of spermatogenesis (Kimble and Ward 1988; Ellis and Schedl 2007). The number of sperms is modulated by the duration of spermatogenesis: the longer spermatogenesis, the larger brood size and the later sexual maturation. As the number of sperms is much lower than that of oocytes, it acts as a limiting factor of total brood size, resulting in a tradeoff between total brood size and generation time (Hodgkin and Barnes 1991; Cutter 2004).

Meiotic X-chromosome nondisjunction (X-nondisjunction) rate increases with maternal age (Rose and Baillie 1979; Luo et al. 2010). The increased X-nondisjunction results in the increased number of male progeny, as *C. elegans* has an XX/XO sex-determination system, where worms with two sex chromosomes (XX) develop into hermaphrodites, but worms with single-sex chromosome (XO) develop into males (Hodgkin 1987). Thus, more sperm production in hermaphrodites may result in an enhancement of progeny production in the late reproductive period, subsequently increasing the brood size and male-production rate, although this idea has not been tested. In previous reports, brood size and male-production rates have thoroughly been investigated in wild strains, natural populations, and laboratory mutants (Huang et al. 2004; Barrière and Félix 2005; Teotónio et al. 2006; Hughes et al. 2007; Luo et al. 2009; Félix and Braendle 2010; Luo et al. 2010; Diaz and Viney 2014; Cutter 2015; Frézal and Félix 2015; Zhang et al. 2021). However, these traits have been measured individually rather than collectively in the same experiment. Thus, to understand the relationship between sperm production and X-nondisjunction, their phenotypic values should be simultaneously quantified.

In this study, we used unmated hermaphrodites of 96 wild *C. elegans* strains and measured their natural variations in brood size and male-production rate during reproductive periods as proxies of sperm-production and X-nondisjunction rates, respectively. The majority of the wild strains exhibited a similar reproductive timing pattern as reported earlier (Klass 1977; Huang et al. 2004; Zhang et al. 2021), and the two traits were not correlated. Our findings imply that although more self-sperm production may increase progeny-production in the late reproductive period, the quality of meiosis could be maintained in the period. We also conducted genome-wide association mapping to identify candidate genetic variants for these phenotypic variations. Our study provides further insights into the natural history of *C. elegans*.

## Materials and Methods

### *C. elegans* culture

All wild strains were maintained at 20 °C on NGM lite plates seeded with OP50. All strains and their phenotypic values are listed in Table S1.

### Measuring brood size and male-production rate

Four virgin hermaphrodites (L4 stage) of each wild strain were transferred separately to new plates, then transferred again every 12–24 h. Three plates were used on the same day, and each experiment was replicated five times. After about 2 d, the offspring could be distinguished by secondary sex characteristics, such as the shape of the tail tip in males. Time constraints prevented examination of all 96 strains at once, so about 15 wild strains were measured in blocks, with strain CB4856 as an internal control. The 96 wild strains were chosen by considering their genetic diversity (Cook et al. 2017). The brood size was calculated as the ratio of the number of progeny to the number of parents, and the male-production rate was calculated as the ratio of the number of males to the number of progeny. These phenotypic values were measured for each period (early, late, and total) and each replicate. Phenotypic change was calculated as (late-early)/(late+early), with the average of all replicates for each strain in the late and early positions. Since NIC260 was an outlier for the male-production rate and ECA36 was an outlier for both the male-production rate and the brood size, these strains were excluded from the following procedures.

### Comparing standard deviations and estimating heritability

To compare the standard deviations of the phenotypic values for each trait and period, tests of equality of standard deviations were conducted using Minitab 19 (https://www.minitab.com). First, the Anderson-Darling test was implemented to test the normality of the distributions of the phenotypic values of brood size and male-production rate. Normality was not rejected for brood size, but was rejected for male-production rate. Therefore, an F-test was performed to compare the standard deviations of brood size, but Bonett’s test and Levene’s test were performed for the standard deviations of the male-production rate. Broad-sense heritability was calculated as described in Gimond *et al*. (2019) using the lmer function in the lme4 package (Bates et al. 2014).

### Analysis of batch effects

Because we divided 96 strains into six groups, we used the CB4856 strains as internal control to test batch effects caused by environmental factors between the groups. To test if the environmental factors made a difference among the six groups, one-way ANOVA was conducted on phenotypic values of the six groups of 15 CB4856 plates. For traits showing differences among the groups (*p*-value <0.05), we conducted multiple comparisons t-test using Bonferroni correction to determine which group pairs were different.

### Correlation analysis and genome-wide association mapping

Correlations were quantified using Pearson’s correlation using DatFrame.corr() from the pandas package in Python (McKinney 2010). Genome-wide association mapping was conducted with the web-based Genetic Mapping in CeNDR (Cook et al. 2017) using the mean phenotypic values of five replicates of 94 wild strains. For phenotypic changes in the early and late periods, the mapping was conducted with the cegwas package, as in the CeNDR webpage (https://github.com/AndersenLab/cegwas2). The variance explained by each QTL and the environmental conditions of the wild strains that we used were obtained from the mapping in CeNDR. JU311 was excluded from the male-production rate change because the total male-production rate of the strain was zero.

### Filtering and categorizing interval variants in associated loci

The interval variants of the male-production rates in the late and total reproductive periods were analyzed only for protein-coding genes in all associated loci. Lists of interval variants were acquired by Genetic Mapping in CeNDR (Cook et al. 2017). These data included the variants’ *p*-values for the phenotypes using Spearman’s correlation tests and their putative impact on the gene, as estimated by SnpEff (Cingolani et al. 2012). The variants of each trait were filtered by *p*-value <0.05, and then among the filtered variants, only those common to both traits were chosen. The common variants were categorized according to their putative impact, and whether they were variants of a gene that is associated with a high incidence of male progeny phenotype. The same process was carried out for genes that had physical, genetic, or regulatory interactions with TGF-β Sma/Mab ligand DLB-1 or insulin/IGF-1 receptor DAF-2 (Harris et al. 2020).

### Calculating linkage disequilibrium between QTL peaks

We measured the linkage disequilibrium between pairs of QTL peaks by calculating the Pearson’s correlation coefficient (Hill and Robertson 1968).

### Data availability statement

All genotypes, marker information, and mapping process are available in the Caenorhabditis elegans Natural Diversity Resource (https://www.elegansvariation.org/). All phenotypes are listed in Table S1 and the whole procedure was detailed in Materials and Methods.

## Results

To understand natural variation in reproductive timing phenotypes, we measured the brood size and the male-production rate, parameters that reflect sperm production and X-nondisjunction rates, respectively. We counted all progeny of 96 wild strains, and checked their sexes in two different reproductive periods: 0–36 (early) and 36–72 (late) hours after adulthood, as *C. elegans* worms produce around 90% of their progeny during these periods (Figure S1, Table S1). Almost all wild strains exhibited continuous phenotypic distributions for all the traits, but we identified a few outliers (Figure S2). ECA36 had the smallest total brood size, of less than 50, which was one-third of the second lowest brood size. ECA36 and NIC260 were outliers in the rate of male production over the total reproductive period. The male-production rates of ECA36 and NIC260 were around 6% and 0.6%, respectively, which were 30 and 3 times higher than the third highest rate. We excluded these two outliers from further analysis. In addition, as we divided 96 strains into six groups to measure their phenotypes, we tested the batch effects in our data using one-way ANOVA. Male-production rates exhibited no significant difference among groups regardless of reproductive timing, but the brood size did (Figure S3, Table S4).

We then compared the phenotypes in the early and late reproductive periods to identify how parental age affects progeny production and male-production rate in wild strains. We found that the average brood size was 148 in the early period and 96 in the late periods (Figure 1A). The average male-production rate was 0.05% in the early period and 0.12% in the late periods (Figure 1B). We also estimated the broad-sense heritability of these traits. Heritability values of brood size were 36%, 62%, and 66% for early, late, and total reproductive periods, respectively (Figure 1A), and the heritability of the total brood size was similar to the previously reported value (63%) (Zhang et al. 2021). The male-production rate exhibited lower heritability values than those of the brood size (9%, 12%, and 18% for male-production rate of early, late, and total reproductive periods, respectively; Figure 1B). Of the 94 wild strains, 87 (92.5%) laid more progeny, and 82 (87.2%) produced fewer male progeny in the early period than in the late period (Figure 1C, D), suggesting that reproductive timing and male-production patterns in wild strains may be similar to those of the reference strain (Klass 1977; Rose and Baillie 1979; Huang et al. 2004).

**Figure 1.**
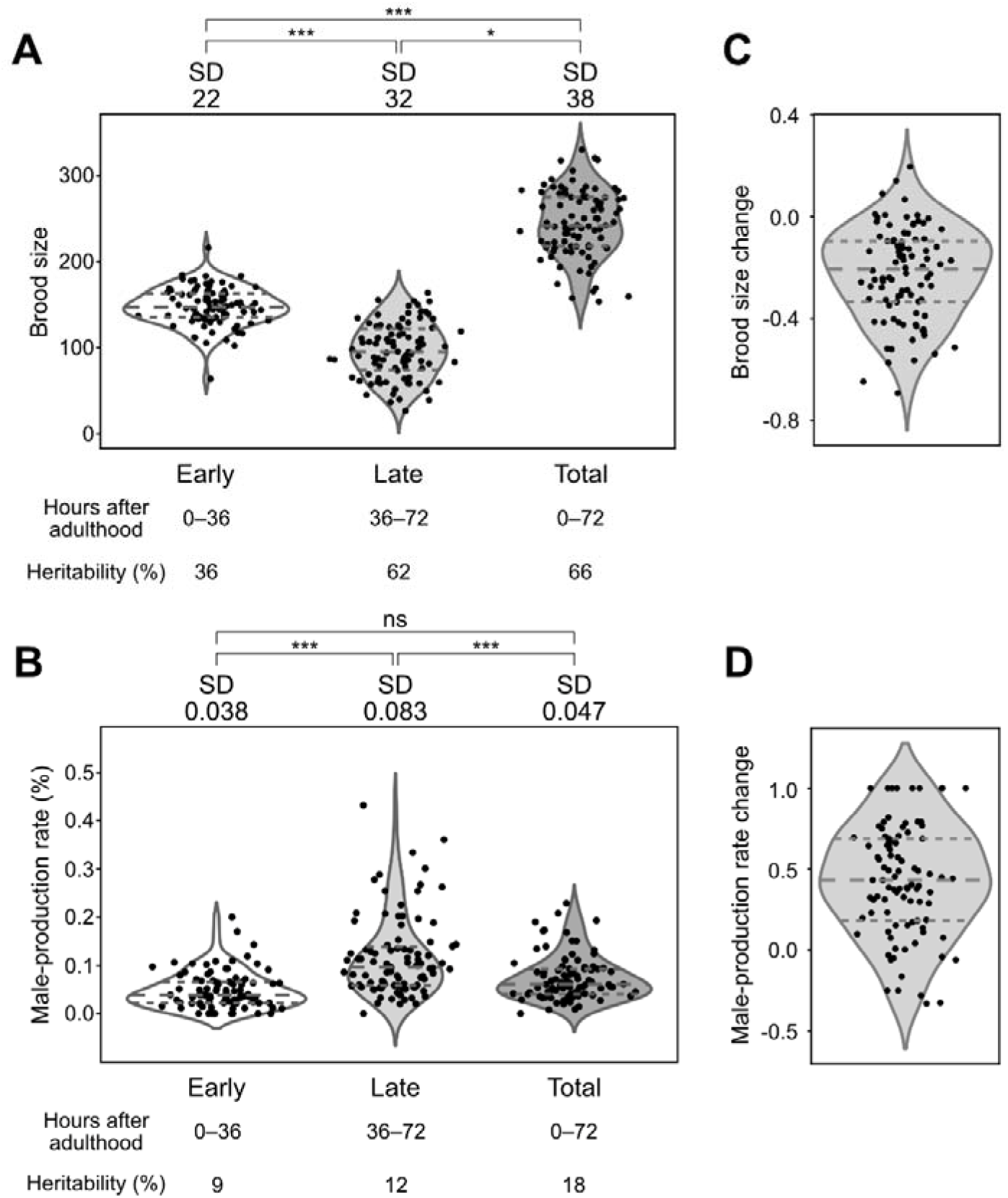
Phenotypic distributions of brood size and male-production rate in each period, and their changes over time. A, B: Distributions of (A) brood size and (B) male-production rate in the early, late and total periods in the 94 wild strains. Each dot represents the average phenotypic value of a wild strain. The *p*-value was used to compare standard deviations using F-tests for brood size, and Bonett’s and Levene’s tests for male-production rate (*: *p* <0.05; ***: *p* <0.001; ns: nonsignificant). Horizontal dashed lines represent (bottom to top) 1st, 2nd and 3rd quartiles. SD, standard deviation C, D: Distributions of change in (C) brood size and (D) male-production rate between two reproductive periods in the 94 wild strains. Phenotypic change was calculated as (late-early)/(late+early).

We tested whether these reproductive timing traits share an underlying genetic architecture by analyzing correlations by genotype for the following eight traits: brood size and male-production rate in early, late and total reproductive periods, and changes in these values between the early and late periods. The brood size and male-production rate were not significantly correlated (Figure 2A), implying that different genetic mechanisms may mediate the two traits. In contrast, the total brood size was weakly positively correlated with the early brood size, but strongly positively correlated with the late brood size, as the phenotypic values of the early brood size were similar among all wild strains (Figure 1A). Phenotypic variations in the late brood size may therefore have primarily affected the total brood size. The total male-production rate was highly positively correlated with the early and late male-production rates, but the early and late male-production rates were weakly correlated to each other, suggesting that they might also have different genetic architectures.

**Figure 2.**
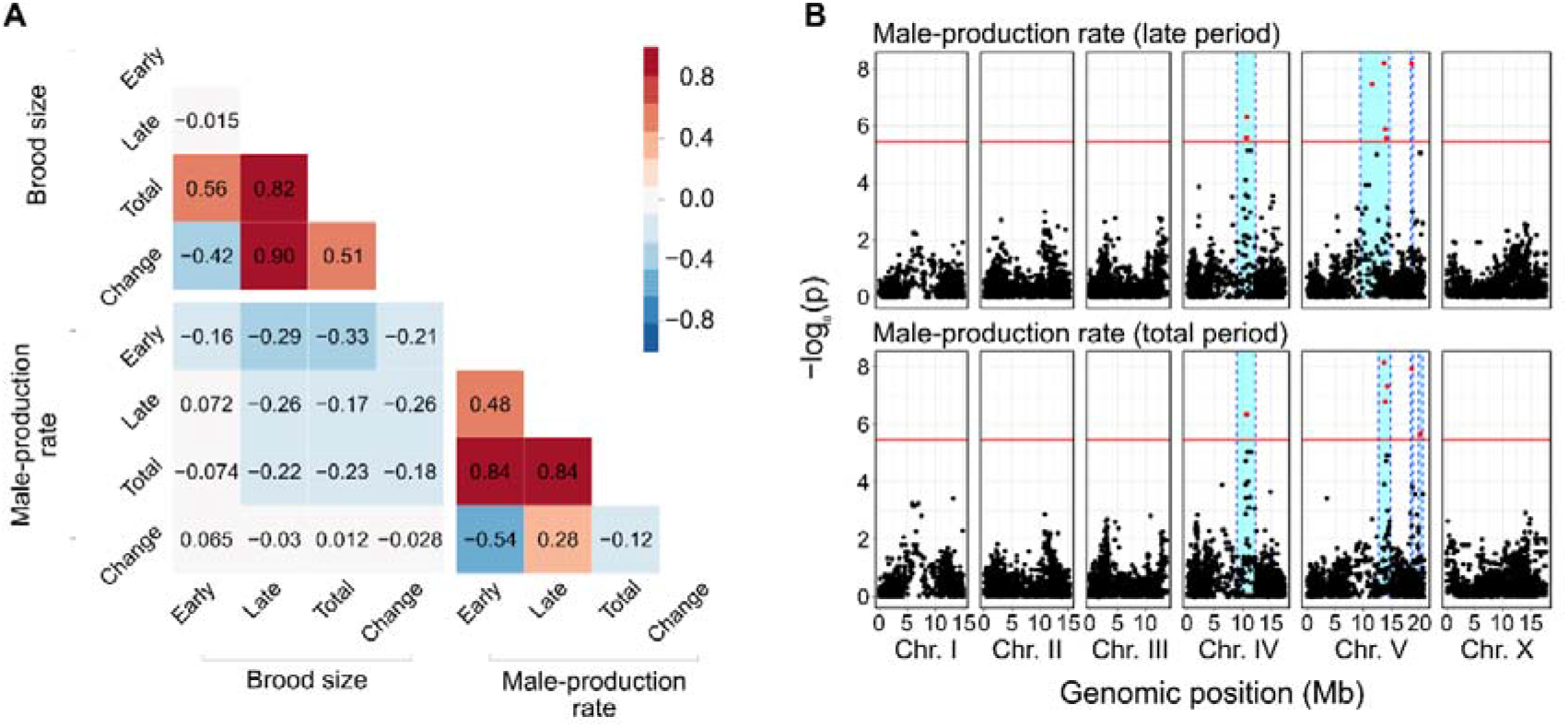
Correlation matrix and genome-wide association mapping for brood size and male-production rate. (A) Heatmap of Pearson correlations between brood size and male-production rate of early, late and total reproductive periods and their changes over time. (B) Manhattan plots for genome-wide association mapping of male-production rates in the late and total periods. Red horizontal lines represent the adjusted Bonferroni cut-off for 5% (-logP= 5.44) and red dots indicate SNVs beyond that threshold. Cyan bars show 95% confidence intervals.

We conducted genome-wide association mapping to investigate the underlying genetic architecture of these traits. We found that several loci were significantly associated with the male-production rates of late and total periods (Figure 2B), but could not find any significant loci for the other six traits (Figure S4). The significant loci for male-production rates in the late and total periods were located on chromosomes IV and V, and largely overlapped, as they were highly correlated (Figure 2A, B), suggesting that they might share significant components of the genetic architecture. We investigated further, using correlations between phenotypic variations and genetic variants at the loci, and found that 3,050 variants were significantly correlated with phenotypes, and were shared between the two traits (Figure 3A).

**Figure 3.**
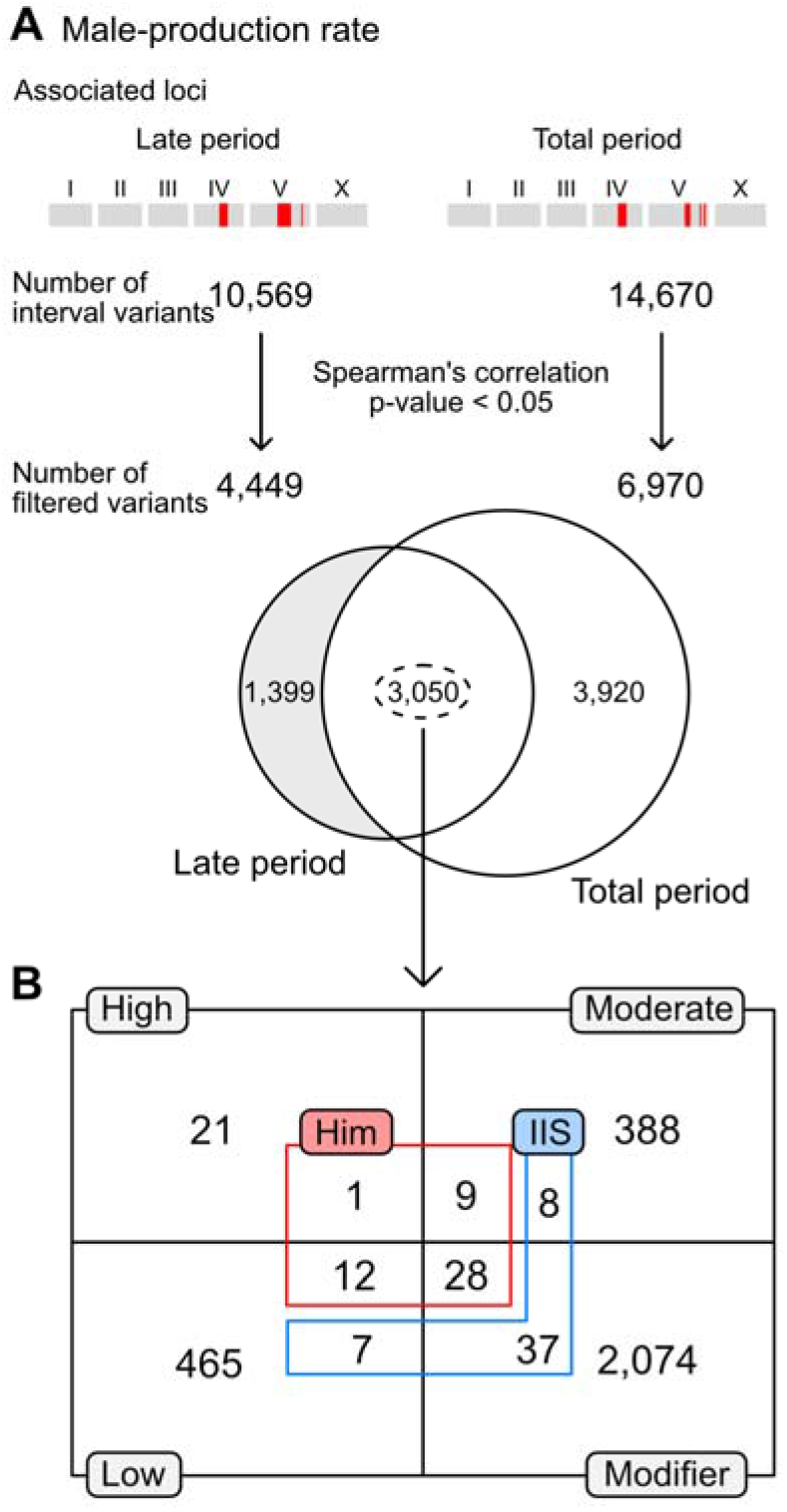
Filtering candidate variants in loci significantly associated with male-production rates in the late and total reproductive periods. (A) Filtering scheme. All interval variants in the loci were filtered according to their correlation with the traits. The loci for the two traits shared thousands of common variants. (B) The number of overlapped and filtered variants were categorized by their putative impacts on genes, as estimated by SnpEff (gray boxes and black lines) (Cingolani et al. 2012). These variants were further categorized by gene functions, including: genes known to interact with the IIS receptor, DAF-2 (blue), or to exhibit Him phenotypes (red), as annotated in the WormBase. Him: high incidence male progeny.

We classified these shared variants according to the known phenotypes that appeared when the functions of the genes containing the variants were impaired by mutation or RNA interference. Among these genes, 11 were known to be associated with the high incidence of male progeny (Him) phenotype (Figure 3B) (Harris et al. 2020). We categorized the shared variants by their putative impacts using SnpEff, and found that *zim-2* contains a high-impact variant and that *coh-3, him-17, him-5, him-8, lex-1, rad-51, sao-1, srgp-1, zim-1, zim-2* and *zim-3* contain moderate-impact, low-impact or modifier variants (Table S2) (Cingolani et al. 2012; Cook et al. 2017).

In addition to genes with the Him phenotype, we analyzed genes that exhibited physical, genetic, or regulatory interactions with major components of the TGF-β or insulin/IGF-1 signaling pathways, which are known to be associated with an increase in the male-production rate with parental age (Luo et al. 2010; Harris et al. 2020). We found some genes that interact with DAF-2, the insulin/IGF-1 receptor (Figure 3B), but could not find any genes for DBL-1, the Sma/Mab TGF-β-related ligand. Among the DAF-2 interacting genes, *daf-10, rpn-7, hsp-12.6, elo-1, fat-4, fat-3, gst-4, daf-14, epi-1, mep-1, let-60, par-5, let-653, fkb-4, mes-4, mtl-2* and *cyp-42A1* have moderate-impact, low-impact or modifier variants (Table S3) (Cingolani et al. 2012; Cook et al. 2017).

We divided wild strains according to their alleles, and calculated the phenotypic variances that could be explained by each locus (Figure 4). We found that for every locus, only five to eight wild strains contained high-male-production rate alleles, and the other approximately 90 strains had low-male-production rate alleles (Figure 4A, B). The variances explained by each locus ranged from 11% to 17% for late male-production rates, and 15%–19% for total male-production rates, values which are similar (Figure 4A, B), and are also similar to the heritability values for those traits (12% and 18%, respectively; Figure 1B). These loci exhibited strong linkage disequilibrium, even between loci on two different chromosomes (Figure 4C, D).

**Figure 4.**
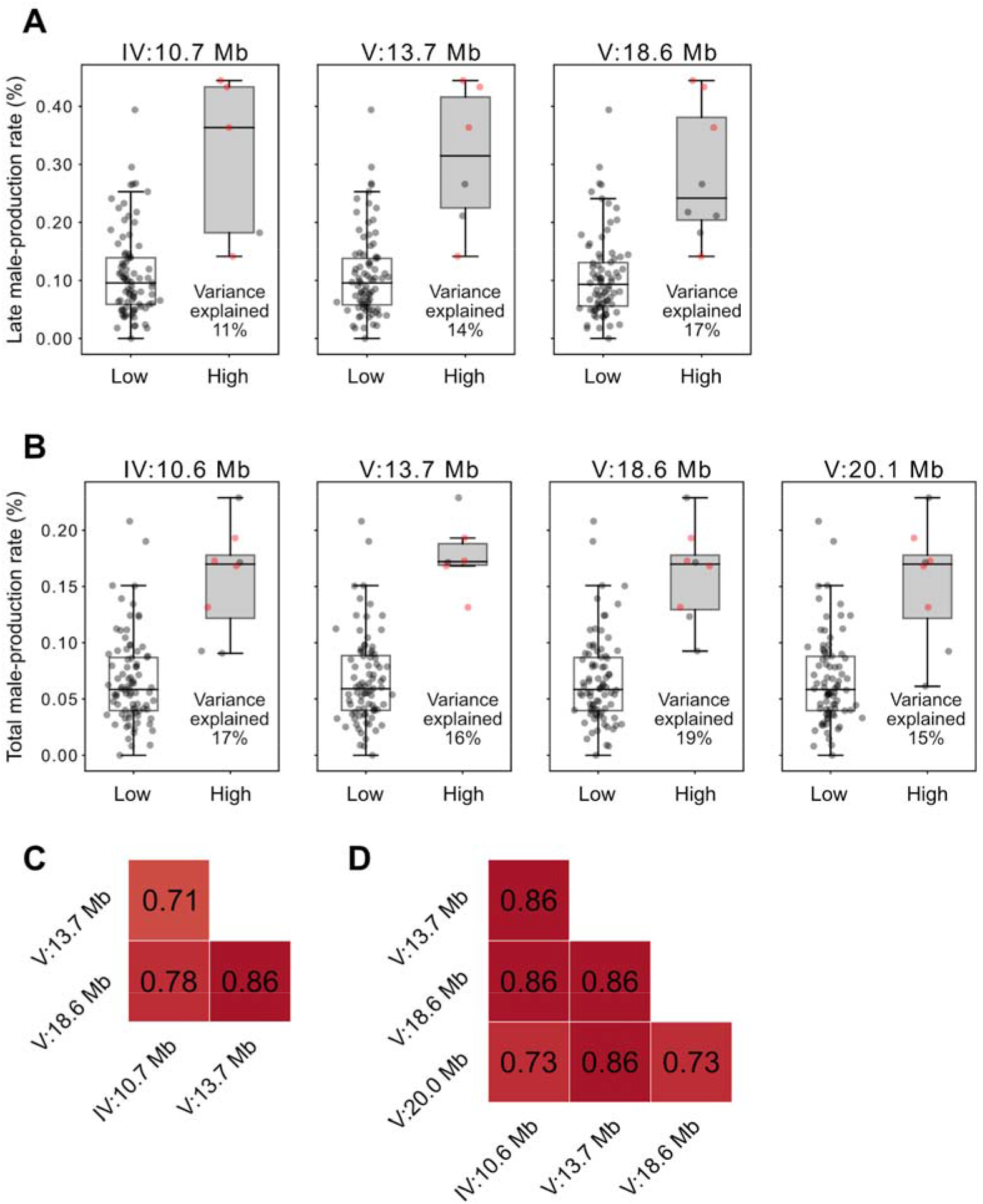
Phenotypic effects of each allele and linkage disequilibrium between associated loci. A, B: Phenotypic differences by peak genotypes of each associated locus for male-production rate in the late (A) and total (B) reproductive periods. Positions of each peak are on the top. Each dot represents the phenotypic value of a strain. The four strains that are marked with red dots had high alleles at all peaks for both the late and total reproductive periods. Variance explained by each locus is represented at the bottom right corner of each box. C, D: Heatmaps of linkage disequilibrium (Pearson’s correlation coefficient, r) between peaks for male-production rate in the late (C) and total (D) reproductive periods.

## Discussion

The reproductive timing in *C. elegans* is associated with sperm production, generation time, and X-nondisjunction. As *C. elegans* is a sequential hermaphrodite, longer sperm-production may delay the generation time and also increase the X-nondisjunction rate owing to the fact that oocyte quality control mechanisms become dysfunctional with age. Intriguingly, we found no significant correlation between progeny number and male-production rate, which were proxies of sperm-production and X-nondisjunction rate, respectively. Most strains produced more progeny and males in the late reproductive period, but strains with larger brood sizes did not always exhibit a higher male-production rate. Furthermore, the male-production rate remained low throughout not only the early but also the total period, implying that spontaneous male production and chromosome segregation were restricted to reduce the high cost of males and aneuploidy (Smith 1978; Cutter et al. 2019). There were two outliers with several or tens of times higher male-production rates in *C. elegans* wild strains, and one of these outliers, ECA36, had a very small brood size. How this outlier survived in the natural environment and which genetic components underlie these extreme phenotypic variations remain elusive.

Different genetic variations may underlie phenotypic variations in sperm production and X-nondisjunction rates, as these traits were only weakly correlated to each other. TGF-β signaling, insulin/IGF-1 signaling, dietary restriction and other pathways modulate reproductive aging in *C. elegans*. TGF-β signaling and insulin/IGF-1 signaling pathways have been reported to be associated with oocyte quality control mechanisms, such as chromosome segregation (Luo et al. 2009; Luo et al. 2010). Notch signaling pathway may affect the oocyte-production rate, as it controls the fates of germline stem cells (Kocsisova et al. 2019). These signaling pathways have genetic variations in wild strains, and they may affect phenotypic variations in reproductive aging phenotypes, but we could not find any loci significantly associated with any of the traits except late and total male-production rates. These loci contain some components of the insulin/IGF-1 pathway and genes known to be related to the X-nondisjunction rate. The genes and variants which mediate the phenotypic differences are still poorly understood.

Our genome-wide association mapping experiments identified only a few associated loci for two out of eight traits. These associated loci contain over 10,000 variants, and such low resolution, in addition to our negative results for other traits, could arise because the number of wild strains that we used was not large enough for fine mapping. It is also possible that the wild strains that we used do not have enough genetic diversity, as *C. elegans* has experienced a chromosome-scale selective sweep, and its hermaphroditism prevents the mixture of different genetic backgrounds (Barrière and Félix 2005; Dolgin et al. 2007; Andersen et al. 2012; Gimond et al. 2013). These characteristics of *C. elegans* might result in low genetic diversity in wild strains. Our data showed that more than 90% of the wild strains that we used contained the same allele at each peak of the associated locus (Figure 4A, B). A recent collection of *C. elegans* from Hawaii exhibits much higher genetic diversity than the set we used, which could possibly be used to resolve other loci associated with the remaining traits (Crombie et al. 2019).

There were several limitations in our experimental procedure. In the previous study, spontaneous male production may have skewed the natural variation in X-nondisjunction rate as mixed worms were possibly used (Teotónio et al. 2006). To exclude this possibility, we used virgin, last larval-stage hermaphrodites. However, they require ∼12 h to reach adulthood; therefore, a maximum 12-h difference in their age may skew the precise estimation of reproductive timing. Specifically, if sperm production timing delays the onset of laying eggs, early and late periods would not be precisely divided into strains with larger brood size. We also did not consider the embryonic lethality and larval arrest, which possibly skews brood size or male-production rate. Moreover, our experimental procedure to maintain and select worms for experiments also varied as population density and maternal age affect nutrient availability and brood size. Wild strains have a CB4856-type *npr-1* allele, and it reduces brood size under high population density (Andersen et al. 2014). Moreover, 1-day adulthood hermaphrodites produce offspring with smaller brood size than that of 3-day adulthood hermaphrodites because older mothers can devote more resources to their eggs (Perez et al. 2017). We did not consider these various sources as we maintained worms with high density (∼100 worms/plate) and selected them regardless of maternal ages. As a result, we may have found that the brood size of our internal control strain, CB4856, was significantly variable among experimental groups (Figure S3).

Our result demonstrates that reproductive timing is an important factor of brood size and male-production rate in *C. elegans* wild strains and that different genetic mechanisms may modulate the two traits. Our associated loci and variants for the male-production rate may act as candidate genetic variants for the trait. It could help in understanding the natural history of *C. elegans*

## Acknowledgements

The Caenorhabditis Genetics Center kindly provided CB4856, and the Caenorhabditis elegans Natural Diversity Resource kindly provided the other strains. We selected genes used to filter QTLs with the help of Wormbase. This work was supported by the Samsung Science and Technology Foundation under Project Number SSTF-BA1501-52. J.Kim was supported by National Research Foundation of Korea grant funded by the Korean government (MEST) [2019R1A6A1A10073437]. J.Lim was supported by a scholarship for basic research, Seoul National University, Seoul, Korea. The authors do not have any conflict of interest to declare. Author Contributions: Conceptualization, J.Kim, J.Lim, and J.Lee; Methodology, J.Lim and J.Kim; Formal Analysis, J.Lim and J.Kim; Investigation, J.Lim and J.Kim; Writing-Original Draft, J.Kim and J.Lim; Writing-Review & Editing, J.Kim, J.Lim and J.Lee; Funding Acquisition, J.Lee; Supervision, J.Lee.

## Supplemental Materials

**Figure S1.**
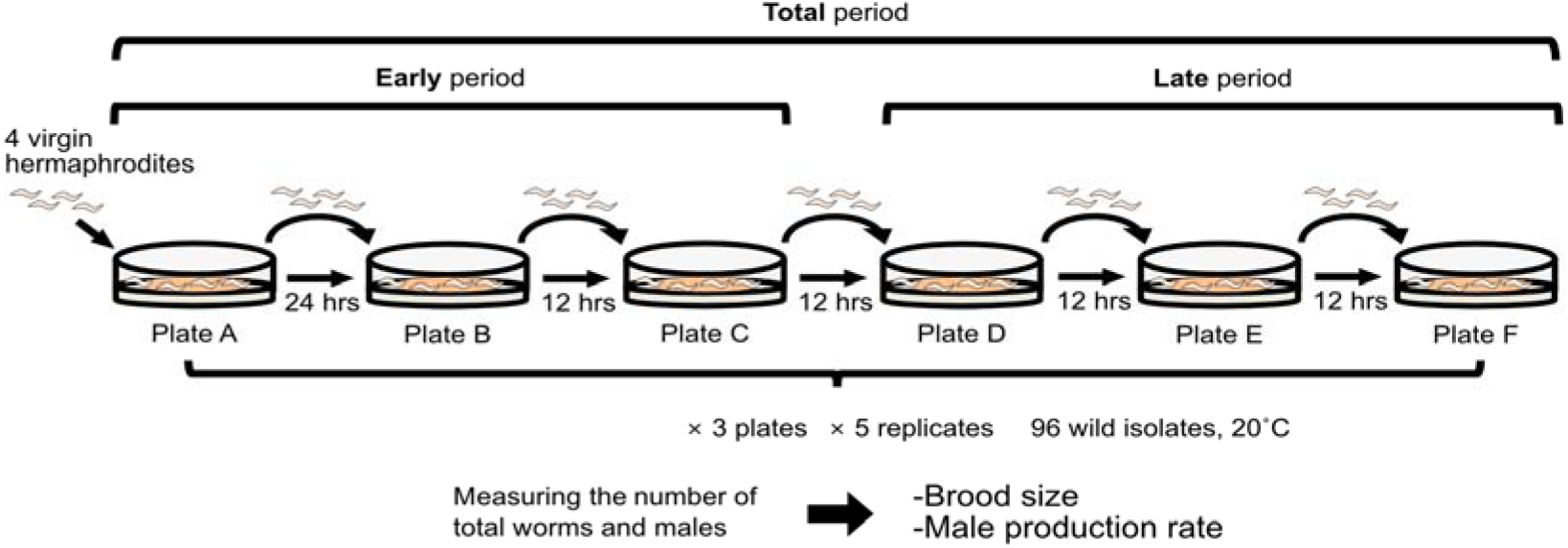
Experimental scheme for measuring the brood size and male-production rate over time. The oocyte-production rates and X-nondisjunction rates were measured in two different reproductive periods. The earlier period was 48 hours after the last larval stage and the later period was the following 36 hours. Four unmated L4 hermaphrodites for each wild strain were raised on the first medium for 24 hours and transferred to fresh media every 12 hours for a total of 84 hours. As the last larval stage of C. *elegans* takes around 12 hours to develop into the adult stage, we used the first 48 hours to 36 hours after entering adulthood. We used three plates for each replicate and counted five replicates in total.

**Figure S2.**
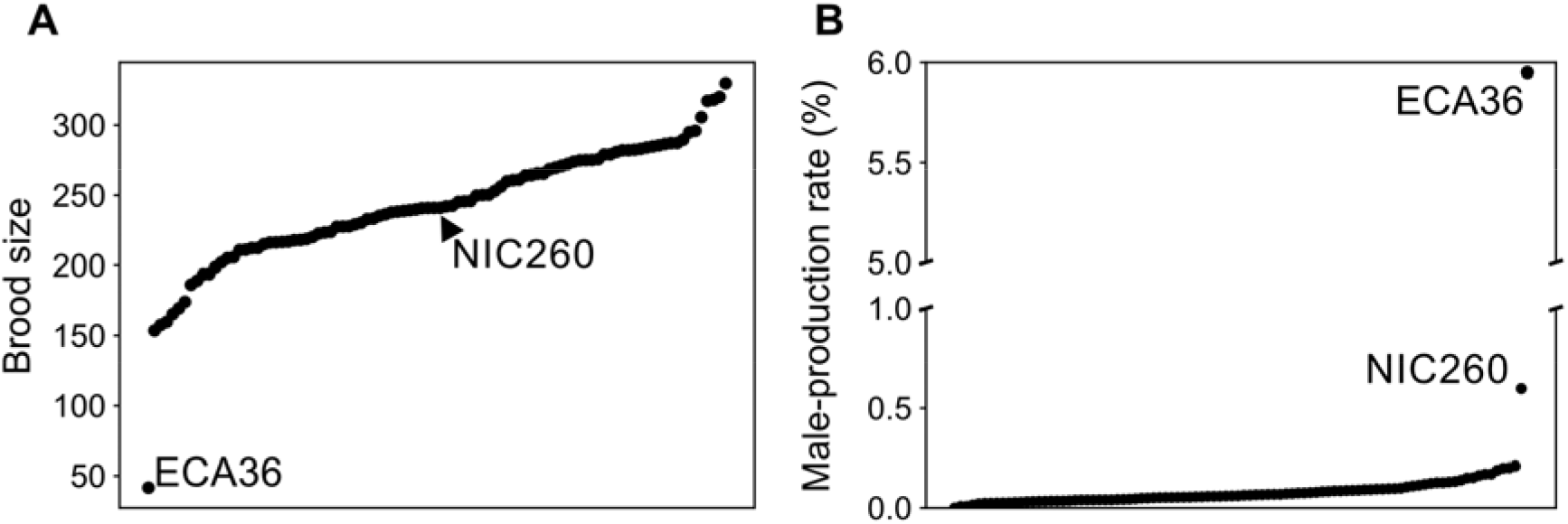
Phenotypic distributions of brood sizes and male-production rates, including outliers. Phenotypic variations for (A) brood size and (B) male-production rate in the total reproductive period including two outliers, ECA36 and NIC260 (arrowheads).

**Figure S3.**
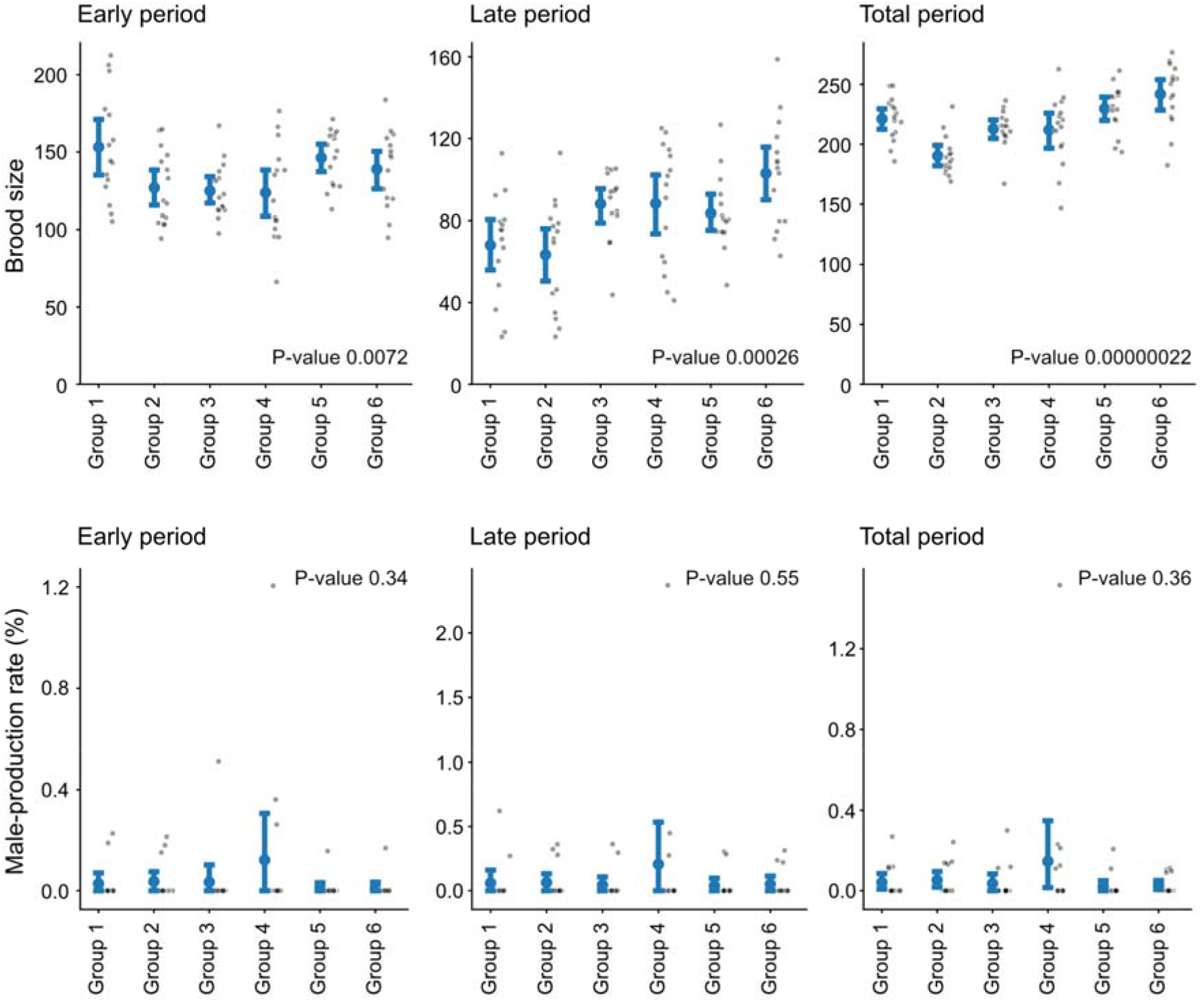
Batch effects in brood size and male-production rate. The dots and scatter plots indicate phenotypic variations among groups of CB4856, our internal control strain. Gray dots represent 15 replicates of each group, and six groups in total were compared. Blue dots indicate averages of replicates in each group; error bars, 95% confidence intervals. *P*-values were analyzed using one-way ANOVA.

**Figure S4.**
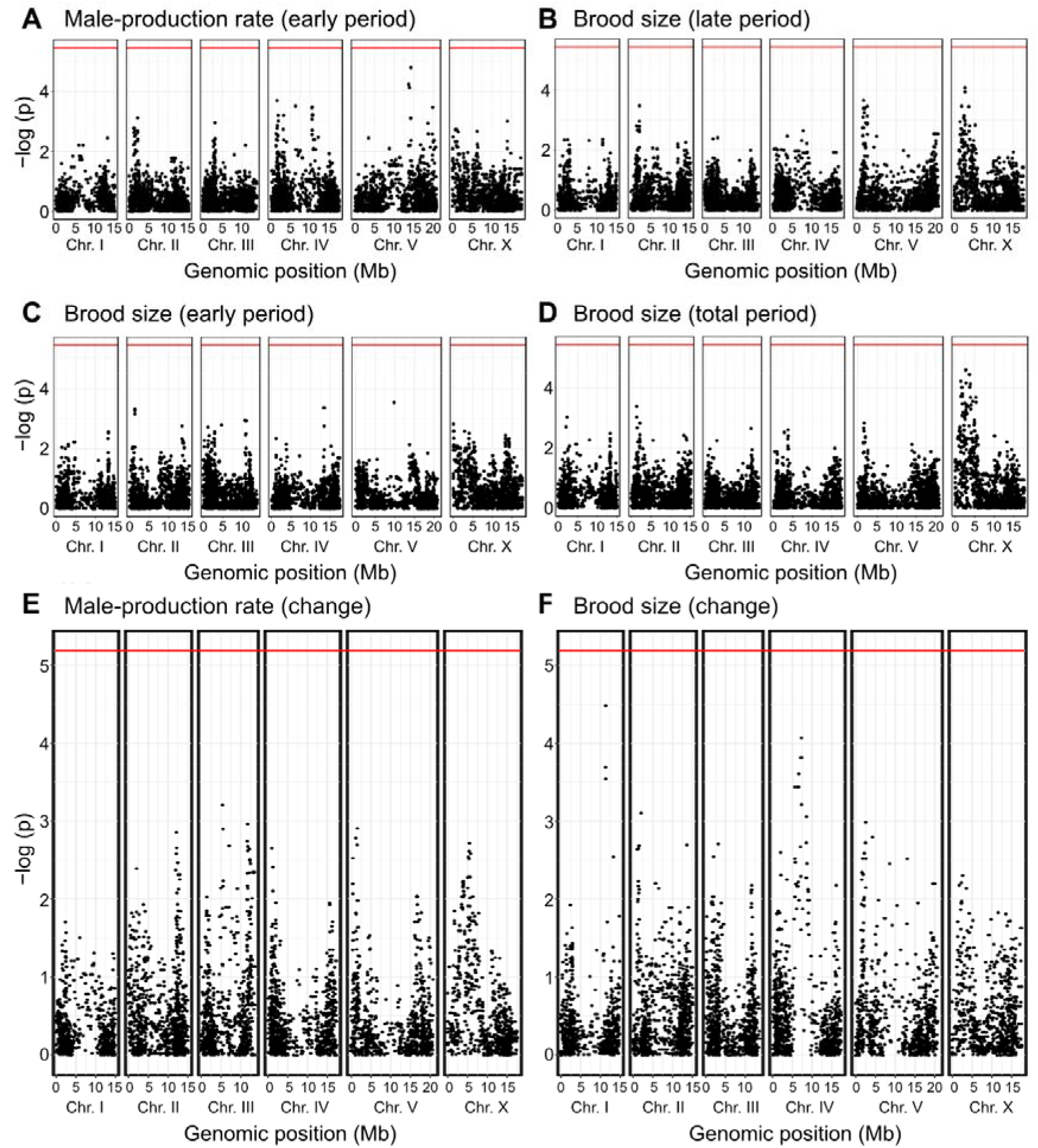
Manhattan plots representing genome-wide association mapping results. (A) male-production rate of the early reproductive period. (B-D) Brood size of late (B), early (C), and total (D) reproductive periods. (E-F) Phenotypic changes between early and late reproductive periods of male-production rate (E) and brood size (F).

(Separately provided)

**Table S1.** Phenotypic values of brood size and male-production rate (%) over time for all wild isolates used in the project

**Table S2.** Candidate variants within genes which exhibit the Him phenotypes

**Table S3.** Candidate variants within genes which interact with the IIS receptor, DAF-2

**Table S4.** Independent samples t-test results between different experimental groups of the CB4856 strain

